# ATLAS: Graph-based 3D RNA Motif Library Incorporating non-Watson-Crick Interactions

**DOI:** 10.64898/2026.02.17.706471

**Authors:** Jingyi Li, Jian Wang, Srinivasan Ekambaram, Nikolay V. Dokholyan

**Affiliations:** Department of Neuroscience & Experimental Therapeutics, Penn State College of Medicine, Hershey, Pennsylvania, USA; Department of Engineering Science and Mechanics, Penn State College of Engineering, University Park, Pennsylvania, USA; Departments of Neurology, University of Virginia, School of Medicine, Charlottesville, Virginia, USA; Depart of Neuroscience, University of Virginia, School of Medicine, Charlottesville, Virginia, USA; Depart of Pharmacology, University of Virginia, School of Medicine, Charlottesville, Virginia, USA; Department of Biomedical Engineering, University of Virginia, School of Medicine, Charlottesville, Virginia, USA; Department of Microbiology, Immunology, & Cancer Biology, University of Virginia, School of Medicine, Charlottesville, Virginia, USA

## Abstract

The structures of RNA exhibit recurrent patterns, which are defined as motifs. Motifs are crucial for the biological functions of RNA molecules. The 3D RNA motif library includes 3D RNA structures, providing a resource for motif-based RNA 3D structure prediction and design. We built the Advanced Template Library for Assembly and Structure (ATLAS) that features graph representations of RNA structures, atomic 3D structures from the Protein Data Bank (PDB), and an isomorphism graph searching algorithm. ATLAS includes nucleotide-level graph representations of RNA motifs and corresponding PDB IDs from the PDB and 3D atomic structures retrieved from the PDB. We provide the web service to search for and download RNA motifs and user-defined structures, supporting non-WC interactions. Graph representations include two types of graphs: motifs with non-Watson-Crick (non-WC) and without non-WC. Compared to existing motif libraries, ATLAS includes graph representations with non-Watson-Crick interactions and pseudoknots and integrates the latest RNA structures from the PDB. We also develop a measure of similarity between RNAs based on shared motifs. We proposed a physics-inspired, time-dependent mechanistic model for RNA evolution, whose steady-state solution fits the global distribution of RNA structural similarity.

## Introduction

The structural complexity of RNA molecules enables their diverse biological functions. Harnessing this complexity in biomedical applications allows for a wide range of uses, including RNA-based therapeutics (1–4) and molecular design. The effectiveness of RNA-based applications greatly depends on the precise configuration of RNA tertiary structures (5, 6). Therefore, designing RNA molecules with specific tertiary structures and functions is critically important. Recurrent structural patterns, known as RNA motifs, are essential for the architecture and function of RNA molecules because these motifs produce physical interactions that lead to functional RNA structures. RNA motifs provide structural frameworks and specific patterns that help RNA molecules perform their biological roles (7). Recently, many methods have been developed for 2D RNA structure predictions (8– 10) and 3D RNA structure predictions (11–29), including discrete molecular dynamics simulation approaches (24–28). Among these methods, motif-based or fragment-based approaches have become powerful tools for predicting RNA tertiary structure (11, 12). Building a comprehensive RNA motif library is a vital step toward advancing RNA assembly, improving design, and deepening our understanding of RNA structure and function.

To address the need for RNA motif libraries, several have been developed. Sarver, Michael et al. developed FR3D (30) for identifying RNA motifs from PDB (31) structures and building motif libraries. Parlea, Lorena G. et al. created the 3D RNA Motif Atlas (32), which includes internal loops, hairpin loops, and three-way junctions with non-WC interactions represented by Varna 2D graphs (33). Tamura, Makio, et al. developed SCOR with further classification based on function (34). Laing, Christian et al. created RNAJunction, focusing on junctions and storing over 12,000 junctions from the PDB (35). Chojnowski, Grzegorz et al. developed RNA Bricks, RNA substructures clustered by similarity using root-mean-square deviation (RMSD) (36). Reinharz et al. designed a graph-based algorithm to extract recurrent interaction networks and build the RNA subunit library (37). However, there remains a need for a single platform that combines a comprehensive motif library—including pseudoknots and non-WC pairs—with detailed graph representations and tools for user-defined structural searches. Compared to 3D atomic models of RNA structures, graph-based representations operate at the residue or motif level, providing simplified views. A graph-based tool (38, 39) has been developed for RNA secondary (40) and tertiary structure prediction (41), and has also been successful in ncRNA-protein interaction prediction (42). We chose a graph representation for motif searching and building, developing an Advanced Template Library for Assembly and Structure (ATLAS). ATLAS is an RNA motif library that includes internal loops, hairpin loops, bulges, pseudoknots, and multiway junctions up to eight-way junctions. It offers three representations: 3D atomic coordinates from PDB files, and nucleotide-level graph models with and without non-WC interactions. By integrating recent PDB data, ATLAS provides a database with over 400,000 motif structures.

## Materials and Methods

### Obtain RNA structures

Obtaining the RNA structures is the first process for building ATLAS (Figure 1). To efficiently obtain the RNA structure data, we developed an automated RNA processing pipeline, RNA downloader. The RNA downloader uses the PDB’s application programming interface (API) to retrieve RNA PDB files that contain the atomic coordinates of these molecules. We built and applied a filter to select the RNA structures and complexes that include RNA structures based on nucleotide types. Some chains also include mutated nucleotides. The filter found the RNA structures by searching for structures that contain A or U or G, or C nucleotides. We identified 8791 RNA structures and RNA-related complexes. The RNA downloader employs format handling, supporting both PDB and mmCIF file formats. The RNA downloader corrects the atom numbering overflow issues and standardizes two-letter chain identifiers to a single-letter format. For the structures obtained from NMR, we utilized only the first model to reduce complexity. This preprocessing pipeline produces standardized RNA coordinate files optimized for subsequent graph-based structural construction, forming the foundation for graph-based RNA analysis.

**Figure 1.**
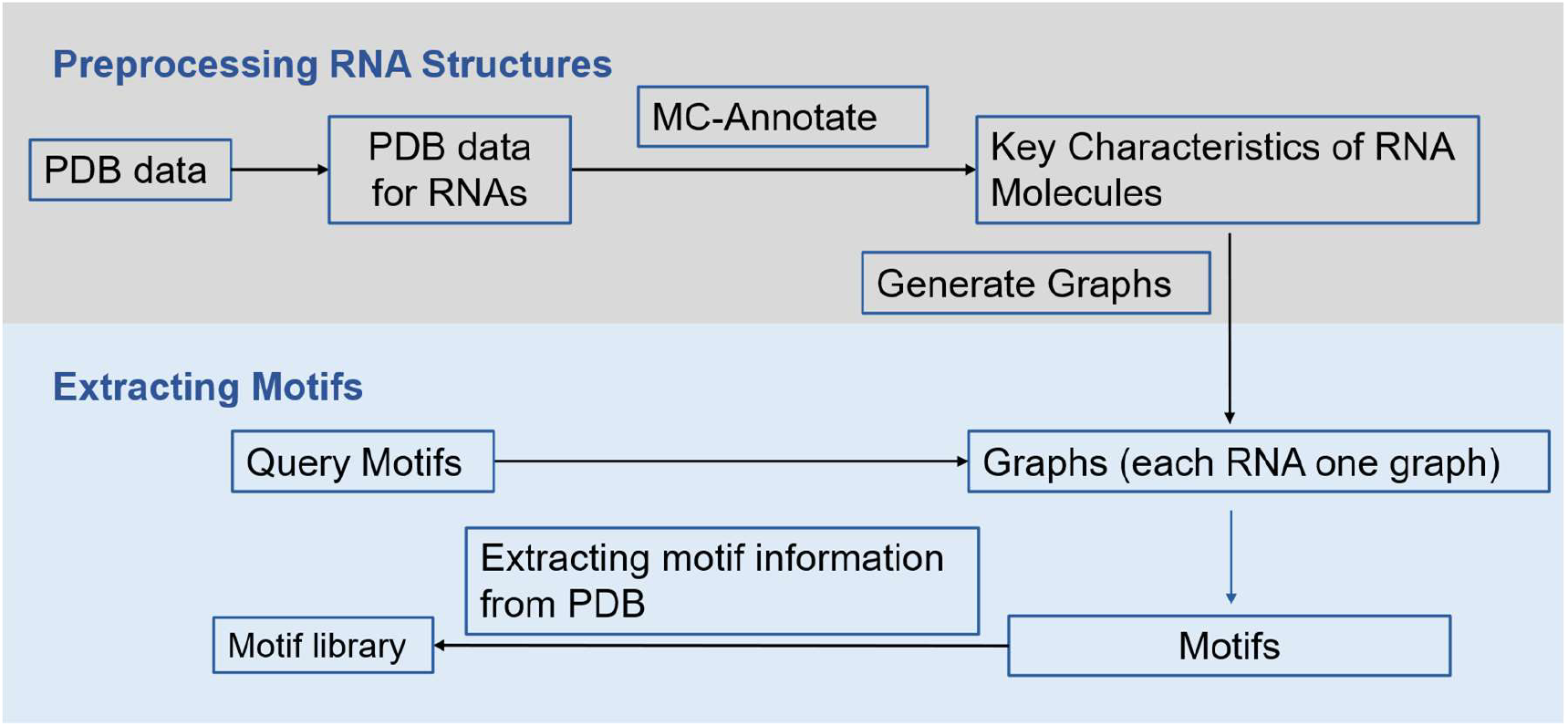
Workflow for constructing ATLAS. The construction of ATLAS includes preprocessing RNA structures and extracting motifs. Preprocessing RNA structures is the process of obtaining the graph representation of RNA structures, and extracting motifs is the process of building a motif library, ATLAS. The construction of ATLAS begins with retrieving RNA structural data from the PDB, which provides 3D atomic structures of biological molecules, including RNA. MC-Annotate(43) is then employed to extract key structural features of RNA molecules. Using this information, graph representations of RNA structures are generated. These graphs are subsequently analyzed to identify RNA motifs. The identified RNA motifs with graph representation and retrieved atomic 3D structures are saved to the motif library, ATLAS.

### Build the graph representing RNA structures

There are different options for representations of RNA structures, including 3D atomic coordinates, dot-bracket notation, and graph-based methods. We chose the nucleotide-level graph-based method because it captures the nucleotide-level RNA geometry and topological properties and represents non-WC interactions. The graph-based method simplifies the representation compared with the 3D atomic structures, and it includes nucleotide-level details compared to motif-level graph representation, in which motifs are represented as nodes, and helices are represented as edges. Nucleotide conformations and base-base interactions act as nodes and edges in the RNA graph. MC-Annotate (version 1.6.2) (43) is used to analyze nucleotide conformations and base-base interactions. MC-Annotate processes RNA structures by analyzing all nucleotide and interaction information, focusing on interactions between nucleotides. It is utilized to identify and categorize these interactions into three types: covalent bonds, WC interactions, and non-WC interactions to simplify the representation. We use one-hot encoding to represent those three types of interactions, assigning different vectors as attributes of edges to differentiate the interaction type (Figure 2a). Additionally, there may also be more than one type of interaction between two nucleotides.

**Figure 2.**
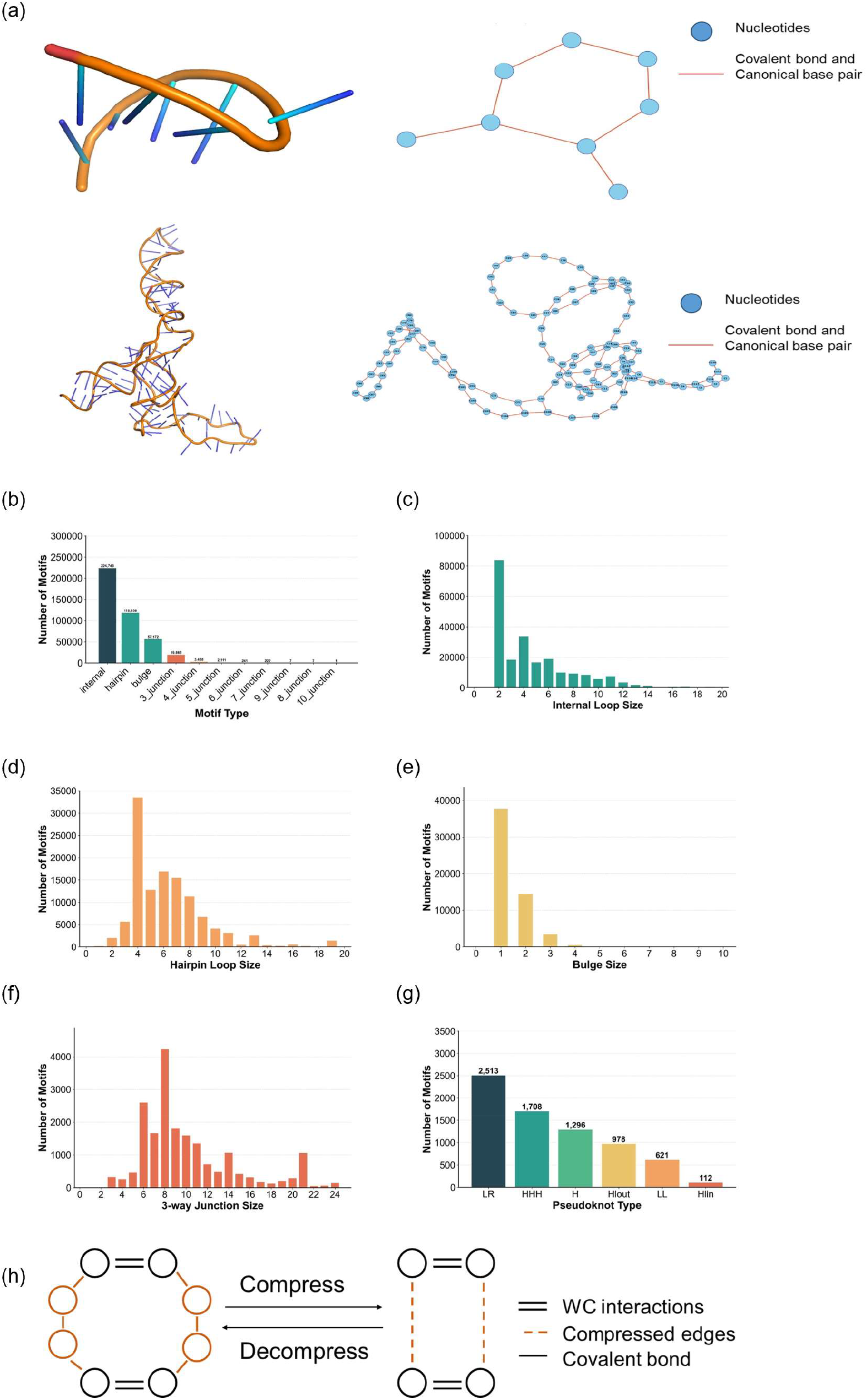
3D structures and their corresponding graph-based representations for RNA molecules as well as motif distributions. (a) The 3D structure of PDB entry 1oj8 (top left), and its graph-based representation (top right). The 3D structure of PDB entry 1c2x (bottom left) and its graph-based representation (bottom right). (b) to (g) shows the motif distributions in ATLAS. (b) Distribution of motif types across the library. (c–g) Size distributions of specific motif categories, including internal loops (c), hairpin loops (d), bulges (e), pseudoknots (f), and 3-way junctions (f). (g) shows the distribution of six types of pseudoknots in ATLAS. LR type and H type have a total of 2513 and 1708 occurrences. HHH type with 1296 occurrences and Hlout type with 978 occurrences, represent intermediate levels of structural complexity and are less common than LR and H type. On the other hand, Hlin with 621 occurrences, and LL with 112 occurrences are relatively rare. (h) shows graph compression and the graph decompression algorithm by compressing successive unpaired nucleotides.

### Graph-based motif search and the construction of the database

We have built graph representations of RNA structures, and the graph representation is utilized for motif searching. To identify motifs in the given RNA database, we utilize the subgraph isomorphism searching algorithm (44). The subgraph searching algorithm searches for predefined graphs in the larger target graphs. We employed this algorithm to identify RNA motifs in graph representations of RNA structures. To simplify the definition of RNA motifs, we do not include non-WC interactions in the search process. The output nucleotide label from the subgraph searching algorithm is used to obtain the corresponding 3D atomic structures and graph representation with and without non-WC interactions. The unique motif ID, the corresponding PDB ID, motif type, 3D atomic coordinates, and graph representations are saved into the SQL database.

### Graph patterns and graph compression for simplifying graphs

Defining the specific query graphs representing the motif structures is essential for subgraph searching. We included only WC base pairs for the RNA motifs, which are defined by WC base pairs, including the hairpin loops, internal loops, bulges, and multiway junctions. We included non-WC base pairs for user-defined searching in the web interface. However, listing all the graphs that represent each type of motif will lead to a significant number of combinations. The graph patterns represent hairpin loops that need to be enumerated with the unpaired nucleotide. We chose one to twenty because high nucleotide numbers lead to thermodynamic instability. The algorithm is built to search through all the defined graph patterns that represent hairpin loops. The subgraph isomorphism searching problem is an NP-complete problem, and the computational cost high and nearly exponentially increases with the graph size (45). It took over an hour on the PC with an i9 13900k CPU to obtain the hairpin loops and store the structures in the database. The enumeration of the hairpin loop contains one number, which is the unpaired nucleotide number. While the internal loop and junctions lead to more complex situations, the specific enumeration of complex motifs needs more graph patterns and a larger graph size than hairpin loops. The increase in graph patterns and size leads to an exponentially increasing computational time. To address these challenges, we developed a graph compression algorithm to decrease both the number of patterns to represent the motif and the size of graph patterns.

The compression algorithm reduces the number of graph patterns and the size of the graph by converting the successive unpaired nodes into edges. The labels of the converted nodes are saved as weights on edges (Figure 2h). The edges are assigned to different attributes, revealing that the edges are special edges storing nodes. The graph-based motif patterns are defined as corresponding to the compressed graphs, reducing the number of motif patterns by avoiding the enumeration of unpaired nucleotides. The subgraph isomorphism searching algorithm operates on the compressed graphs and defined RNA patterns. The output graphs are then decompressed to obtain the original structures of the RNA motif, subtracting for RNA structure. The decompression algorithm reduces the complexity of RNA and motif patterns, accelerating the overall motif searching.

### Pseudoknots detection

RNA pseudoknots are complex motifs frequently found in viral and cellular RNAs. Pseudoknots are formed by hairpin loops with additional base pairing between the single-strand region and the unpaired region of hairpin motifs (46). Pseudoknots play an important role in gene expression and regulation (47). We included the pseudoknots in ATLAS because pseudoknots are an essential part of the RNA motif library. However, pseudoknots do not have a single graph pattern, which makes it hard to define the motif pattern. To address this challenge, we used the dot-bracket representation to represent the RNA structures. The graph representations of RNAs without non-WC interactions were used to convert to the dot-bracket representation to maintain consistency in RNA annotations. We built an algorithm to label the sequence with dot-bracket form with dots, brackets, and square brackets. Dot-bracket representation employs the dot and the pairwise bracket to represent the unpaired and paired nucleotides. For more complex structures, like pseudoknots, extended dot-bracket notation is used. The search algorithm finds pseudoknots by searching the string for subsequences enclosed by square brackets. Then the sequences within square brackets are extracted and extended when unmatched brackets are encountered to ensure complete annotation of pseudoknot regions. Finally, the 3D atomic coordinates, graph, and dot-bracket representations for the pseudoknots were saved in the SQLite database.

### Web interface

We developed a web interface for public access to ATLAS. The site is implemented in Python/Flask and serves data from a SQLite database. The backend exposes routes for the homepage and search workflow, a custom motif workflow, and downloads. For standard search, the user submits motif type and size via POST. The server filters ATLAS.db by motif class and size and returns rows. Results are rendered as an HTML table and exported to CSV. The CSV includes the ID, motif type, PDB ID, and nucleotide number columns from the database. PDB files that meet the filter are found and packed into downloadable ZIP files. For custom search, the frontend provides a canvas where users define a query graph by adding nucleotides and selecting edge types, including covalent, Watson–Crick, and non–Watson–Crick. The graph is serialized to JSON and passed to an external attribute-constrained subgraph isomorphism routine. Matches are written to a results database, and the site then reads the database and provides the download options, including CSV and ZIP. The frontend is implemented with vanilla HTML/CSS/JavaScript, and Flask handles routing, templating, and file streaming.

## Results

### The database web interface

ATLAS web interface (http://atlas.dokhlab.org/) has a motif searching block, a custom searching block, and a presentation of distribution block. In the searching part, with the chosen motif type and size, all the outputs are shown, and all the results can be downloaded either in a CSV file or a Zip file, including all the PDBs for the motifs. The custom searching block provides a canvas for users to draw the structure with nucleotides and interactions, supporting non-WC interactions. The user-defined structures are utilized to search for RNA substructures among the PDB database. Also, the user can download the results in a CSV file or a zip file with PDB files. The distribution part shows the motif type distribution and the distributions for different types of motifs. Users can use the website’s search tool or the custom search tool to access the database.

### Distribution of motifs

ATLAS is an RNA motif library that includes five types of RNA motifs: internal loops, hairpin loops, bulges, multiway junctions, and pseudoknots. Analysis of the library reveals the distribution of those five motif types and sizes within the current PDB. ATLAS incorporates atomic 3D coordinates and graph representations with or without non-WC interactions and includes 433,996 motifs. Internal loops are the most frequent motif type, representing 51.8% of the identified motifs. The percentages of hairpin loops, bulges, 3-way junctions, and pseudoknots are 27.4%, 13.2%, 4.6%, and 1.7% respectively. The size of the motif is defined as the number of unpaired nucleotides. Internal loops contain two base pairs and two sets of unpaired nucleotides, and the minimal nucleotide number of internal loops is 2, which is a 1-1 internal loop (Figure 2c). 97.4% of the internal loops range from 2 to 12 nucleotides, and the occurrence of internal loops of size 2 (2 unpaired nucleotides) is 84,182 (37.5%). The size distribution of hairpins shows that 89.3% of hairpin loops range from 2 to 10 nucleotides. The most frequent size of hairpin loops is 4, and the occurrence is 33,555 (28.2%). Bulges have a relatively narrow size distribution, with most of them ranging from 1 to 3 nucleotides (98.0%). The most frequent size is 1 (66.2%). Most of the sizes of 3-way junctions are from 5 to 14 (81.0%). The most frequent size of 3-way junctions is 8 (21.4%).

### Pseudoknots Distribution

ATLAS captures the pseudoknots, and 7,228 pseudoknots were found and then further classified into six groups: Long Range (LR), HHH, H, Hlout, LL and Hlin. Pseudoknots are defined as two helical regions with a connection(48). All six types of pseudoknots were defined and found (Figure 3a-f). LR Type is the most frequent type of pseudoknots, which is 34.8%. The high frequency is because it includes a diverse range of pseudoknots, making it the most inclusive type of pseudoknot. Following the LR type, the HHH type (23.6%) is identified as the second most common pseudoknot, also called the kissing hairpin, and it is formed by the additional interactions between unpaired parts of two hairpin loops. The H type (17.9%) is the third most common. As the simplest structure among all types of pseudoknots, the H type is also the simplest possible structure, which likely contributes to its frequency. Hlout (13.5%) has one more hairpin loop at the single-strand region. Those types with less frequency, including LL and Hlin, are likely associated with their more complex structural conformations (Figure 2g). The percentages of LL and Hlin are 8.6% and 1.5%, respectively.

**Figure 3.**
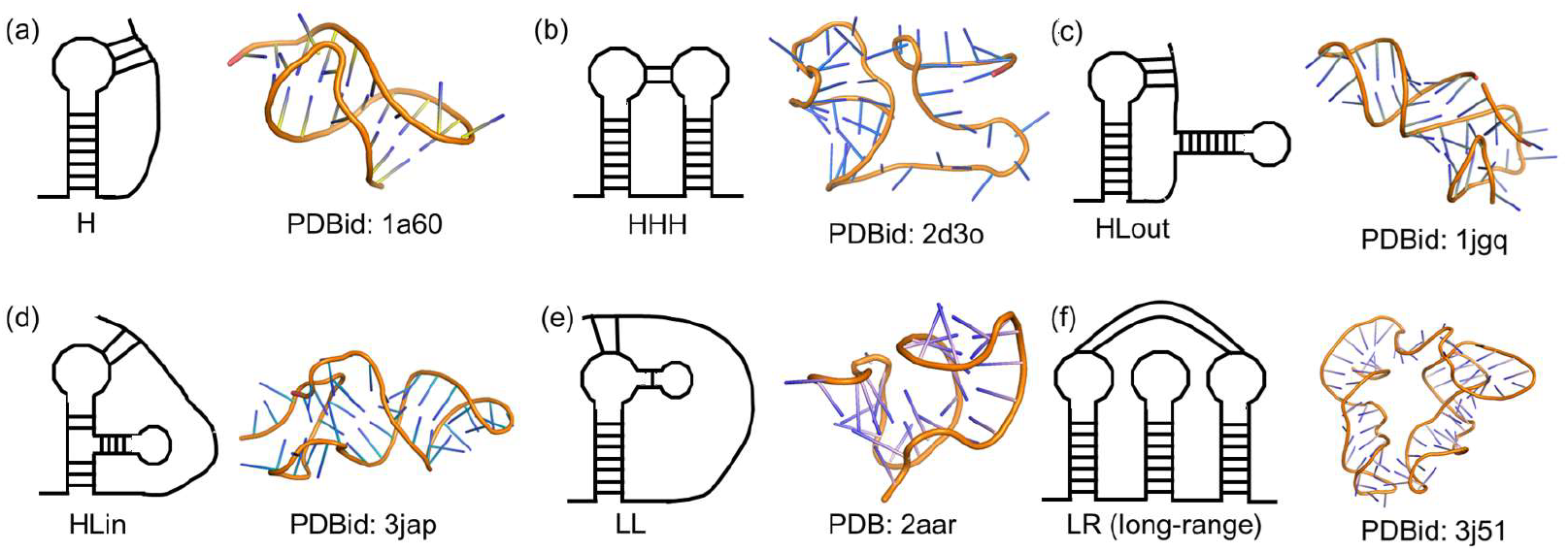
Type of pseudoknots and corresponding 3D structures, as well as the distributions of pseudoknots. Pseudoknots are categorized into six structural clusters: 1H Type (a), HHH Type(b), Hlout Type (c), Hlin Type (d), LL Type (e), and RL Type (f). Structures not fitting other categories are classified as LR Type, which includes diverse and complex pseudoknots with over 2500 nucleotides.

### Classification of motifs with non-WC interactions

#### Algorithm 1

RNA Motif Classification Algorithm

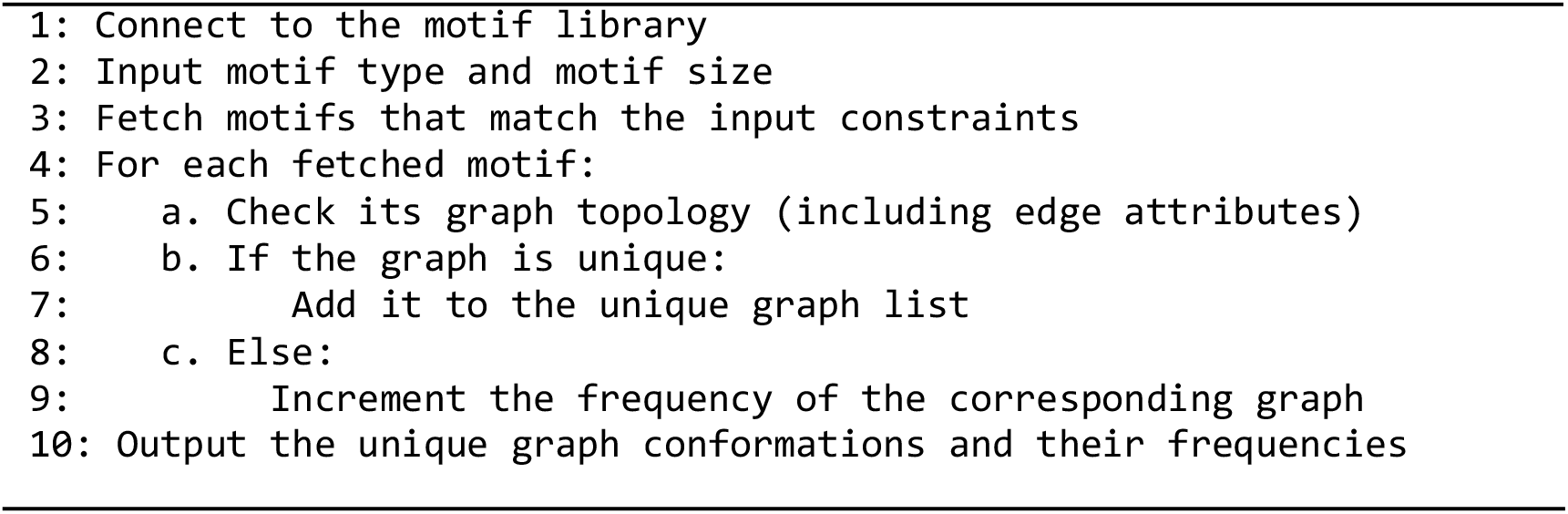

ATLAS incorporates non-WC interactions, which form the basis of RNA structure and contribute to the formation of RNA motifs. Motifs with identical WC-based classifications exhibit distinct structures when non-WC interactions are considered essential to the RNA structures (49). ATLAS incorporates non-WC interactions, enabling a refined classification of motifs. We developed an algorithm for the automatic classification of motifs with non-WC interaction based on their graph representations. The algorithm finds all the graph-based motif structures. RNA motifs are classified automatically based on graph topologies. Motifs are considered distinct based on their type, size, and the nature of their base-pairing interactions, encompassing both Watson-Crick and non-Watson-Crick pairs. Non-WC interactions significantly increase the diversity of RNA motifs. For example, hairpin loops with 4 unpaired nucleotides exhibit 1487 unique conformations. The largest cluster of hairpin loops contains hairpin loops with two non-WC base pairs between the unpaired nucleotides (Figure 4a). For all hairpin loops, the top three frequent motifs are from hairpin loops with six nucleotides. The most frequent conformations for hairpin loops with 5, 7, and 8 nucleotides are the conformations without any non-WC interactions (Figure 4c). The corresponding distributions (Figure 4b, d, f) illustrate the frequencies of these conformations. The 1-1 internal loop with 6 nucleotides is the most frequent internal loop.

**Figure 4.**
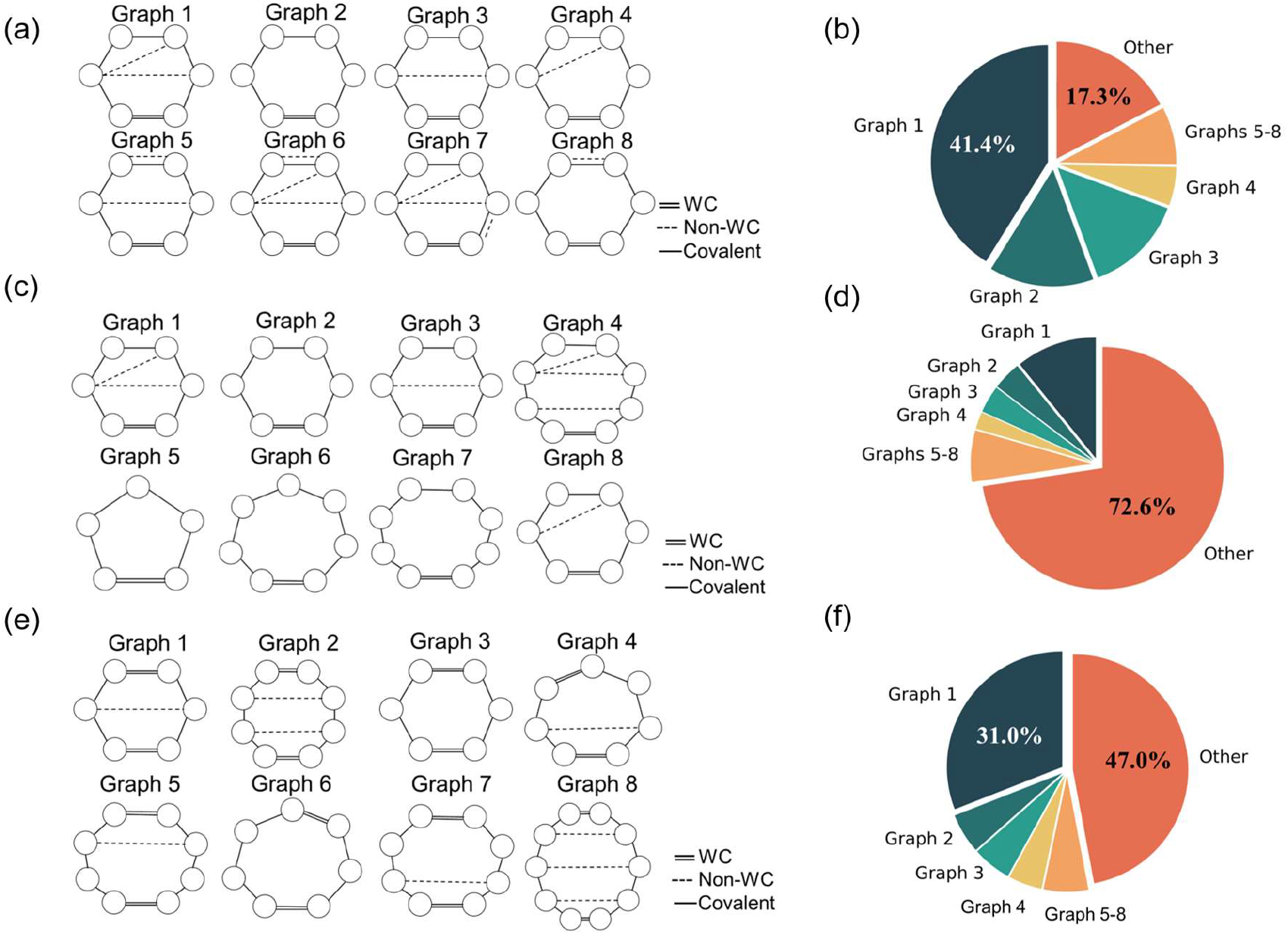
Motif conformations for motifs considering non-WC interactions and corresponding distributions. (a) presents the 12 most frequent hairpin loops as shown. The graph with the highest frequency is labeled Graph 1. The pie chart (b) illustrates the distribution of graphs for hairpin loops with 6 nucleotides, with graph 1 taking 41.4%. (c) illustrates the top 8 conformations for all hairpin loops. (d) shows the distributions for the conformations, revealing the diversity of hairpin loops. (e) and (f) present the top 8 frequent internal loops and distribution repetitively, revealing that 1-1 internal loops are the most frequent, which is 31.0% of all internal loops.

### Motif-based RNA similarity

ATLAS includes the graph representations of motifs. RNA structure is related to RNA function, yet sequence-based and structure-based comparisons often fail to capture the subtle and complex features of RNA folding. We developed a motif-based RNA similarity algorithm (MBRS) to quantify RNA similarity based on motif similarity using the data from ATLAS. MBRS incorporates multiple RNA motif types, including internal loops, hairpin loops, bulges, and junctions, to calculate the RNA similarity. We obtain graph representations of motifs from ATLAS with different attributes representing the different types of motifs. The graph-based similarity score to quantify the motif-motif similarity *S*_*m*, σ (*m*)_ is calculated by an enhanced WL Kernel Similarity algorithm (50). The similarity is normalized from 0 to 1. Then, the weighted motif-motif similarity is aggregated to obtain RNA similarity,

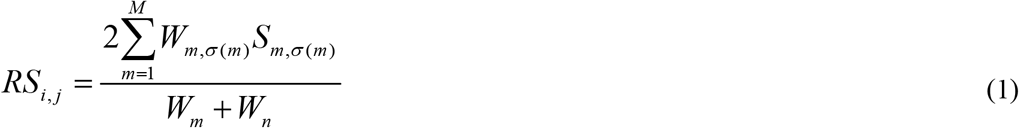

σ(m) is the best match between two sets of motifs to maximize the unweighted similarity, finding the optimized 1-1 mapping between motifs to maximize the RNA similarity. The best match is found with the linear sum assignment algorithm to find the best match. The pairwise motif-motif similarity matrix is first calculated by *S*_*i, j*_ = *Similarity*(*Motif*_*i*_, *Motif* _*j*_), and the cost matrix is then calculated by 1− *S*_*i, j*_.

*W*_*m*_ is the weight calculated with *W*_*m*_ = *L*_*m*_ × (1+ *NC*_*m*_), which *L*_*m*_ is the number of nucleotides in the motif *m*, and *NC*_*m*_ is the number of non-WC base pairs in the motif *m*.

The pairwise RNA similarity distribution is shown in Figure 5a. We hypothesize that RNAs in the same functional family have higher intra-family similarity than inter-family similarity. To test the hypothesis, we computed intra-family similarity scores for RNA pairs within the same functional groups and compared them to randomly sampled inter-family scores from Rfam (51). The violin plot of similarity scores is shown in Figure 5b. The Mann-Whitney U test(52) yielded a p-value less than 0.05. The significant difference between intra-family and inter-family RNA pairs supports our hypothesis that RNA within the same functional family has higher motif-based RNA similarity.

**Figure 5.**
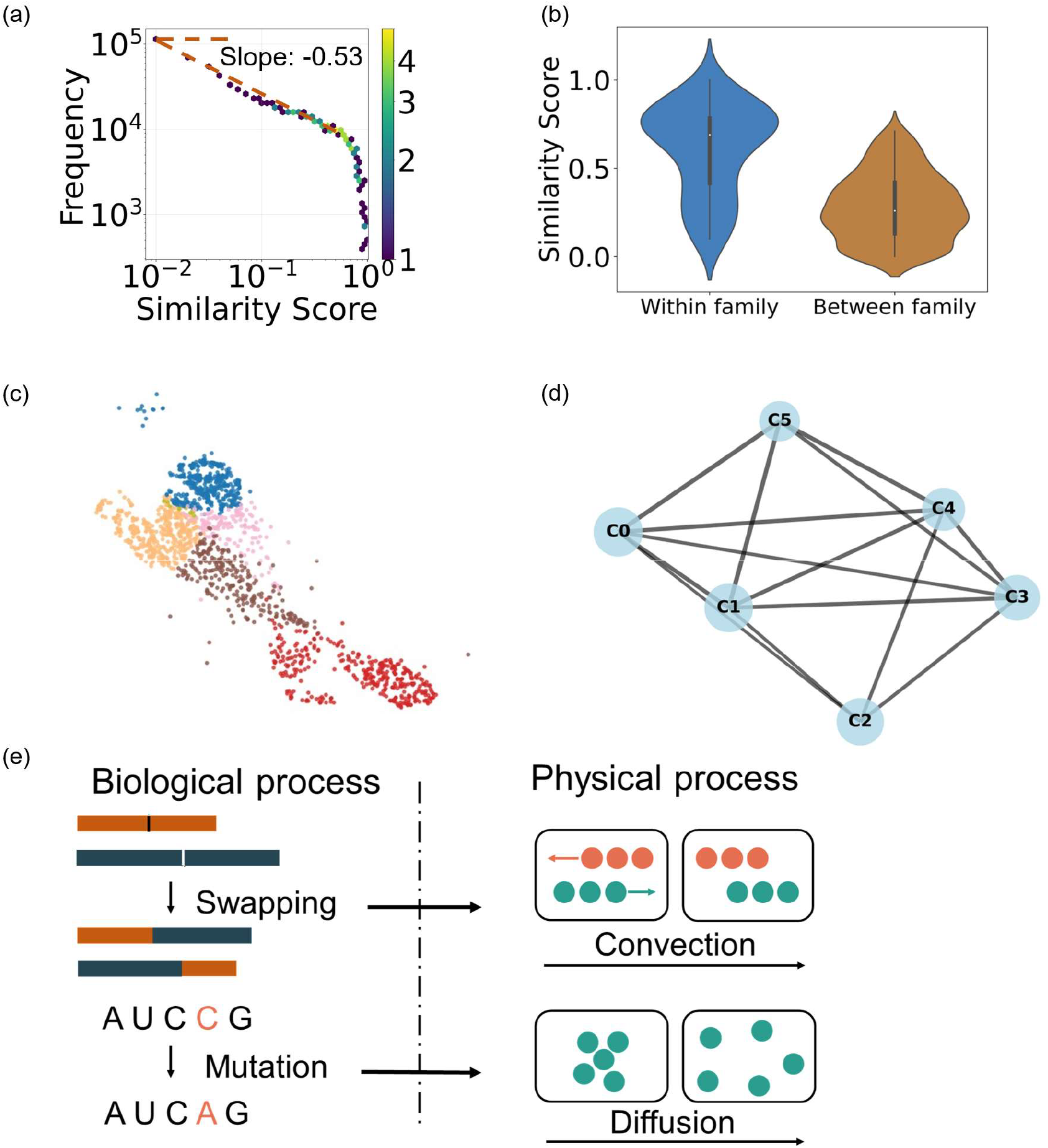
RNA similarity distribution, similarity comparisons, and clustering patterns. (a) presents the distribution of the RNA-RNA similarity score, ranging from 0 to 1, plotted with the logarithmic x-axis and logarithmic y-axis. The color represents the number of data points falling within each hexagonal bin. Violin plot (b) shows the similarity score distributions for the within-family and between-family groups. The within-family group (mean 0.61, median = 0.69) has higher similarity compared to the between-family group (mean = 0.27, median = 0.26). (c) is a scatter plot with dots representing RNAs. The coordinate is based on the distance between RNAs, calculated by the similarities between them. All points are clustered and color-coded based on the similarity score. (d) RNA cluster meta-graph illustrating inter-cluster relationships. Nodes represent RNA clusters, each containing more than 10 RNAs. (e) In RNA evolution, the swapping may correspond to the convection term, and the mutation may correspond to the diffusion term in the Fokker-Planck equation.

According to the pairwise similarity, the Leiden algorithm (53) is used for obtaining the cluster of RNAs. We cluster the RNA, and color the nodes with different colors for different clusters Figure 5c, and represent clusters with super nodes as shown in Figure 5d. Our motif-based RNA similarity map provides a unique perspective to study the evolution of RNA molecules.

### Modeling RNA evolution based on the Fokker-Planck equation

RNA evolution processes contribute to the structural and functional diversity of RNA molecules (54–56). The RNA World hypothesis (57) proposes that early life functioned based on RNA. RNA motifs are recurring 3D building blocks that appear in many RNA molecules. Some motifs, such as hairpin loops, internal loops, bulges, and junctions, are found in many RNAs. Those motifs are related to important biological functions. Because of the evolutionary pressure, they are conserved across RNA molecules (58). During RNA evolution, complementary mutations do not always cause the overall folding change(58). We hypothesize that the distribution of structural similarity between RNA molecules (Figure 5a) is the consequence of evolutionary processes. Here, we propose a simplified model of RNA evolution that explains this distribution.

We found that the frequency *f*_*R*_ is proportional to 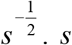 is the similarity score, and *f*_*R*_ is the corresponding frequency, which is the probability density function after normalization. We posit that during evolution, RNA motifs “diffuse” from one structure to another, for example, through various processes, such as mutation. In addition, there may be a “convective drift” of motifs as they may contribute to structural stabilities or function may be caused by crossover (59, 60), whereby two strands of daughter chromatids exchange a segment. (Figure 5e). The Fokker-Planck equation describes the time evolution of the probability density function *p*(*x, t*) of the position:

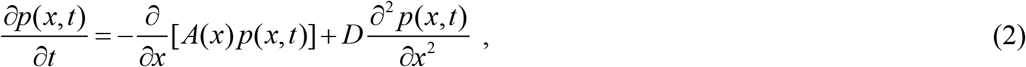

where *A*(*x*) *p*(*x, t*) is the drift term, also called the convection term, which represents a systematic change in the system because of the driving force. 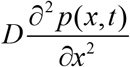 is the diffusion term, which is caused by the random fluctuations. *D* is the diffusion coefficient. In the law of mass conservation in continuous mechanics,

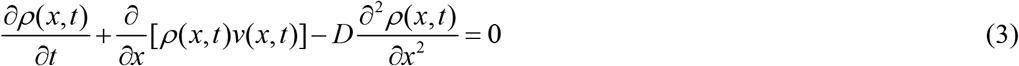

*ρ* (*x, t*) is the mass density, *v*(*x, t*) is the fluid’s velocity field, corresponding to the convection part in the Fokker-Planck equation. 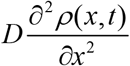 represent the mass moving from high concentration to low concentration.

By physical analogy, we hypothesize that the change in the frequency of RNA pairwise similarity can be described by the equation in the same format as the Fokker-Planck and mass conservation in continuous mechanics.

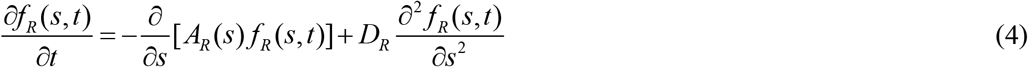

*f*_*R*_ (*s,t*) describes the evolution of frequency over time and similarity. *A*_*R*_ (*s*) *f*_*R*_ (*s, t*) is the convection term that systematically changes the frequency distribution, including the process motif swapping. Motif swapping disrupts the homologous motifs and generates novel combinations, which reduces the similarities. 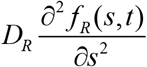 represents the similarity fluctuation caused by mutations. For a steady state, 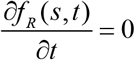, and substitute the equation into equation (4).

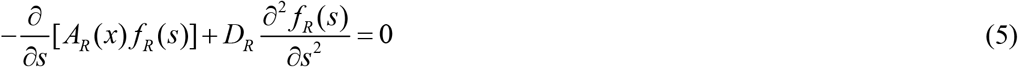

The solution to equation (5) is

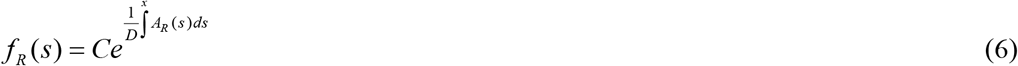

To explain the similarity distribution as shown in Figure 5a:

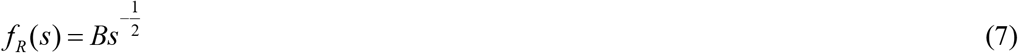

We find the solution, which is the drift coefficient,

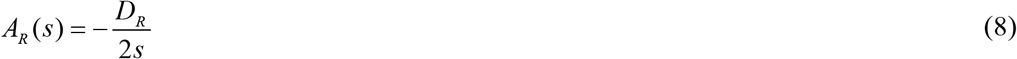

We assume the drift term corresponds to the swapping process, occurring at a constant rate. When two RNAs have a slight similarity, the driving force for changing the similarity is high because the swapping operations are more likely to lead to a higher change in similarity. While for RNA structures with high similarity, it is more likely to have the swapping behavior between motifs or swappable units, with structures with high similarity.

The RNA evolution equation provides a potential explanation for the pairwise RNA similarity distribution.

## Discussion

In recent years, graph-based tools and algorithms have provided novel approaches for studying RNA motifs. The utility of graph-based methods contributes the RNA analysis. RNA structures are represented by graphs at the atomic level, residue level, and coarse-grained level. Graph theory offers a unique analysis method for obtaining insight into RNA and motif structures. However, identifying RNA motifs without explicit definitions has been a challenge in motif-related research. Because of the diversity of the RNA molecules and the complexity of RNA motif structures, manually defining RNA motif structures leads to many combinations. The limitations highlighted the need for approaches that identify the RNA motif with the partial definition of motif structures.

Graph-based representations of RNA structures enable the application of unsupervised learning techniques for identifying RNA classification. The unsupervised learning algorithm can group similar structural patterns without manually explicit definitions of RNA motifs. For complex RNA structures, it is much easier to define the partial motif descriptions without complex details. These partial definitions could be utilized as search templates to extract features of more generalized motifs. The output dataset could be used as input for training Graph Neural Networks (GNNs).

Many RNA motifs, especially complex ones, do not have a single, complete, and formal rule-based definition that can identify every single instance. Instead, the definitions are more like descriptions of common features. ATLAS provides data to train the GNN to search for RNA motifs even without an explicit definition. Our approach incorporates the existing knowledge, which is the partial pattern and the GNN tools, to uncover the hidden pattern for RNA motifs. By learning from ATLAS data, GNN can generalize to identify the full family of motifs. For example, in the case of pseudoknots, we use dot-bracket notation to represent pseudoknots. The parentheses and square brackets are typically sufficient for most cases, but different brackets are needed for representing more complex pseudoknot interactions. Having the completeness of definitions for pseudoknots with complex structures has always been a challenge. By providing GNN partial patterns, for instance, the simple representation of pseudoknots, the model could then learn from the partial patterns and generalize the understanding of RNA motif structures. The generalization ability of the GNN model is used to identify RNA motifs with a simple definition.

Unsupervised learning excels in data clustering by recognizing patterns, relationships, and groupings in data without the need for predefined labels or categories. This approach can be applied to RNA structure datasets to cluster RNA motifs with similar features or substructures. By analyzing features of the structure of RNA graphs, the unsupervised learning method can identify motifs with subtle structural similarities, even if these motifs differ in terms of graph size or interactions, corresponding to the same type of RNA motifs with one or several more nucleotides. These recognized features contribute to classifying RNA motifs into different families. The classification provides important insights into the structural diversity and even functional roles of RNA molecules. This clustering process will also deepen our understanding of RNA structural architecture. By integrating graph-based representations with unsupervised learning techniques, we may be able to overcome the limitations of traditional, sequence-based or descriptive structural-based motif discovery and classification methods. This approach will not only facilitate the discovery of new RNA motifs but also further refine existing motif libraries and greatly contribute to the evolutionary advancement of our understanding of RNA structure and function.

MBRS can be used to predict the function of the newly found RNA. By calculating the similarity between the newly discovered RNA and the RNA with known function, high similarity may lead to a high probability of functional similarity. MBRS provides a way for RNA classification and RNA evolution. By calculating the similarity between RNAs across evolutionary lineages, researchers may find evidence of how motifs are conserved or changed over time.

The model of RNA evolution, based on the Fokker-Planck equation, provides a quantitative possible explanation for the observed steady-state distribution of pairwise RNA structural similarities. Because the Fokker-Planck equation is time-dependent, the RNA evolution model offers a framework to describe the dynamic process of RNA evolution. The RNA evolution model may reveal the underlying time-dependent mechanisms of RNA evolution, offering a tool for research to investigate how RNAs undergo random structural exploration via mutation and oriented change via processes like swapping.

3D RNA design is the process of creating RNA molecules with specific 3D structures with atomic-level accuracy (61). However, RNA folding presents significant diversity because of the structural complexity (62). The single-strand region adopts diverse 3D conformations, resulting in complex folding possibilities. Motif-based RNA design or nucleotide-based RNA design incorporating the knowledge from motif structures reduces the dimension of designing space for the same size of RNA compared with the nucleotide-based design.

3D RNA structure prediction and 3D protein structure prediction share similarities because they build a mapping from the linear space, which is the sequence of RNA, to the 3D space, which is the 3D structure. Given the success of fragment-based computational tools in protein design, such as deep learning algorithms (63) and generative models (64), these computational tools may be adapted for RNA design by modifying them to account for the unique properties of RNA.

ATLAS provides a foundation for 3D RNA prediction and 3D RNA design by providing atomic 3D coordinates and nucleotide-level graph representations of motifs. These representations provide options for developing computation tools. By leveraging ATLAS and a helix library, a reinforcement learning-based algorithm may be developed by choosing optimized building blocks and assembling them. Additionally, motifs in the ATLAS may act as constraints for generative models. Biological constraints may guide the generative process, leading to a biologically meaningful RNA structure. As a database of RNA motif structures, ATLAS serves as a basis for the application of advanced computational tools, including 3D RNA prediction and 3D RNA design.

## Conclusion

We have developed ATLAS, an RNA motif library that includes graph-based and geometric representations of atomic structures extracted from the PDB. ATLAS includes five types of RNA motifs, including hairpin loops, bulges, internal loops, junctions, and pseudoknots. ATLAS provides the residue-level graph structure of RNA motifs and the corresponding RNA PDB IDs with and without non-WC interactions. ATLAS includes more than 400,000 motifs and three representations for each motif. Our analysis shows that incorporating non-WC interactions results in more structural complexity. By representing motif structures with both graph and atomic coordinates, ATLAS offers an optional resource for studying structural diversity, RNA similarity calculation, and RNA evolution. We provide a web interface to search for and download the motif data and user-defined RNA substructures, including non-WC interactions. We developed a novel motif-based RNA similarity method, MBRS, which provides a new direction for calculating RNA similarity by focusing on motif similarity. The RNA similarity calculated by MBRS is related to the function, providing a tool for exploring the RNA functional and evolutionary relationships. We propose a Fokker-Planck-like model to describe the RNA evolution process. The model gives a possible explanation for the distribution of pairwise RNA similarity and its frequency. Although ATLAS is currently limited to known motif types, it establishes a foundation for advancing RNA prediction, design, similarity calculation, and evolution research in the future.

## Data Availability

Source codes and test data are deposited at <u>dokhlab / atlas — Bitbucket</u>, and the website for the database is hosted at http://atlas.dokhlab.org.

## Acknowledgments

We acknowledge support from the National Institutes of Health (R35 GM134864), the National Science Foundation (2040667), and the Passan Foundation. This project was also supported by the Penn State College of Medicine’s Artificial Intelligence and Biomedical Informatics Program.

